# Molecular Investigations of Selected Spike Protein Mutations in SARS-CoV-2: Delta and Omicron Variants and Omicron Subvariants

**DOI:** 10.1101/2022.05.25.493484

**Authors:** Urmi Roy

## Abstract

Among the multiple SARS-CoV-2 variants recently reported, the Delta variant has generated most perilous and widespread effects. Another variant, Omicron, has been identified specifically for its high transmissibility. Omicron contains numerous spike (S) protein mutations and in numbers much larger than those of its predecessor variants. In this report we discuss some essential structural aspects and time-based structure changes of a selected set of spike protein mutations within the Delta and Omicron variants. The expected impact of multiple-point mutations within the spike protein’s receptor-binding domain (RBD) and S1 of these variants are examined. Additionally, RBD of the more recently emerged subvariants BA.4, BA.5 and BA.2.12.1 are discussed. Within the latter group, BA.5 represents globally, the most prevalent form of SARS-CoV-2 at the present time. Temporal mutation profile for the subvariant BF.7 and currently circulating variants of interest (VOI) and variants under monitoring (VUMs) including XBB.1.5, BQ.1, BA.2.75, CH.1.1, XBB and XBF are computationally explored here briefly with the expectation that these structural data will be helpful to identify drug targets and to neutralize antibodies for the evolving variants/subvariants of SARS-CoV-2.

## 1. Introduction

Since WHO declared a Public Health Emergency due to SARS-Cov-2 in January 2020 [1], several different variants of coronavirus have emerged. Among these variants, the Delta strain has been reported to be particularly aggressive due to its fast transmissibility and strong infectivity. The Delta variant, also known as the B.1.617.2 strain, was first identified in India, and was listed among the CDC’s “Variants of Concern” [2]. The highly contagious B.1.1.529 strain, Omicron, was first identified in South Africa in December 2021 [3], and the presence of numerous spike (S) protein mutations in Omicron makes this variant highly transmissible. The BA.2 subvariant of Omicron, containing many additional mutations in its S protein has been a predominant COVID 19 species for several months until recently [4]. At this present time, another subvariant BA.5 largely represents the most active form of coronavirus worldwide. Among the other recent Omicron subvariants, BA.4 and BA.2.12.1 are also notable [5]. BF.7, a sublineage of BA.5, also has been spreading in some countries. Lately, XBB.1.5 is designated as circulating variant of interest (VOI) by WHO on March 15, 2023. Additionally as of March 22, 2023, WHO has listed BQ.1, BA.2.75, CH.1.1, XBB and XBF as circulating variants under monitoring (VUMs). In this report these recently listed VOI and VUMs are briefly mentioned in the Supplementary Materials (SM) [6]. The mutations within the recent SARS-COV-2 variants and subvariants are listed in SM Tables S1-S3. As new variants/sub variants of SARS-CoV-2 continue to emerge, they are likely to be associated with more epitope escape mutations with greater epidemiological fitness.

High performance computational tools have the potential of aiding the ongoing efforts to identify the proper therapeutic targets for these pathogenic variants. In this regard, structure-based computational immunology and bioinformatics based methods of immuno-engineering play a critical role in detecting proper drug targets as well as in designing new vaccines [7-11].

In previous reports, the author has discussed the time-based structures of various immunologically significant proteins and their structure-functions relationships [12-14]. We have also reported the impacts of mutations on the receptor binding domain (RBD) of SARS-CoV-2 [15,16]. Recently we have also examined the impact of Zn ion coordinated Angiotensin II peptides on the spike 1 (S1)-angiotensin converting enzyme-2 (ACE2) receptor binding and the role of these engineered peptides in bio catalytic processes [17]. In the present communication, we will analyze most of the RBD mutations observed in the Delta and Omicron variants and Omicron subvariants BA.2, BA.4/BA.5 and BA2.12.1 and determine their structural changes and stability as functions of time. Additionally, the corresponding cases for BF.7, BQ.1, BA.2.75, XBB, XBB.1.5 and XBF are briefly mentioned in this report. Results for the time based structures of wild type (wt) SARS-CoV-2 S1 and the S1 mutations within the Delta, Omicron, and BA.2 will also be included here.

A Recent report indicates that the Omicron subvariant BA.2 has multiple point mutations similar to those of Omicron variant and several Omicron subvariants, including BA.2 belong to Omicron [2]. However, previously identified BA.2 may contain several different mutations. Since the BA.2 variant described in this report is based on that earlier data, and since SARS-CoV-2 mutations tend to be variable, the data based on older version of BA.2 was not removed. The formerly identified BA.2 subvariant is labelled as BA.2 throughout this report [2]. Subvariants BA.4 and BA.5 contain identical RBD mutations, the details of which are included in tables S2.

Recently a few cases of Deltacron recombinant species are identified in some countries. However, the phylogenetic tree of SARS-CoV-2 Omicron subvariants BA.1, BA.1.1, BA.2, BA.2.12.1 and BA.4/5 do not reflect proximity between the Delta and Omicron variants [18]. The RBD and S1 mutations within the Mu variant are also briefly explored here as well. The Mu was briefly discussed in this report was described as “Variants Being Monitored” by CDC [2]. Two relatively new subvariants BA.2.75 and BF.7 are also included in the SM section in addition to the circulating VOI and VUMs.

## 2. Materials and Methods

### 2.1. Mutation Steps

The spike (S) protein of the SARS CoV-2 comprises two subunits, S1 and S2. This paper is centered on the wt and mutant variants of the RBD and S1 subunit. The X-ray crystal structure 6M0J.PDB was selected for the wt RBD simulations [19]. Starting from the wt RBD structure, the mutant variants were generated using the VMD mutator plugin graphical user window (GUI). Since most of the mutations are within the newly emerged subvariants, and since these types of mutations tend to evolve consistently, they are frequently created in the computational approach. We have analyzed the selected RBD mutations within the B.1.617.2 strain, the Delta variant and the B.1.1.529 strain, the Omicron variant, along with an exploration of the subvariants BA.2, BA.4/BA5 and BA.2.12.1.

The I-TASSER server was used to model the structure of SARS-CoV-2 S1 [20]. The wt S1 was mostly prepared based on the CryoEM structure of 6VYB.PDB; chain B [21]. Due to the lack of experimental data, part of the protein sequences have still been missing in the published literature, making those sequences difficult to model; some of these missing sequences have been based here on the PDB structure 6ZP2 and other resources reported elsewhere [4,22]. In addition to the RBD mutations, wt, Delta, Omicron and BA.2 have been also examined in terms of their S1 characteristics.

The current work is a continuation of our previously published report [16], where the RBD mutations within B.1.1.7, B.1.351, and P1 lineages were discussed; these mutations were initially found in the UK, South Africa, and Brazil. The S1 sub unit of Delta strain based on 6Z97.PDB was also briefly mentioned there [16]. This present communication, discusses the RBD of Omicron and its newly emerged subvariants. In addition, a comparison of the Delta and Omicron S1 is carried out using a larger model structure where most of the missing sequences are restored.

The B.1.621 strain, also known as the Mu variant, as well as the subvariants, BA.2.75, BF.7 and circulating VOI and VUMs are briefly mentioned in the SM. Initially, the BA.4/BA.5 subvariants were considered to have two mutations L452R, F486V and a reversion of Q493R [5]. To examine the implications of this consideration, the RBD of a possible subvariant with two mutations L452R and F486V along with the reversion of Q493R are also discussed in the SM section. These results help to establish a perspective for comparing the earlier description with the latest version of BA.4/BA.5. The current version of the BA.4/BA.5 contains mutations that exist in the newer form of BA.2 in addition to the above two mutations and a reverse mutation. The selected RBD and S1 mutations used in all simulation experiments have been listed in SM Table S1-S3, and some of the mutations have been updated further through the course of this study.

### 2.2. MD Simulations

For the simulation purpose the QwikMD and NAMD packages were used. For the data analyses and visualization VMD program was utilized [23-25]. The implicit solvation system was used with the Generalized Born Solvent-Accessible Surface Area [26]. Initially the PSF structure files were prepared by deleting atoms and renaming residues. Energy minimization was performed for 2000 steps, and annealing was executed for 0.24 ns using a temperature increase from 60K to 300K. Equilibration was performed for 0.04 ns at 300K. After performing the initial minimization, annealing and equilibration steps, the MD production run was procured using the NVT ensemble. In all cases the integration time was selected for 2fs. The CHARMM36 force field was applied for all simulations. For the annealing and equilibration protocols, the backbone atoms were restrained, while no atoms were constrained during the final MD simulation. The detailed experimental procedures have been described elsewhere [15]. The exploratory simulations described in this report are based on the initial conformational samplings that provide a general insight into the proteins’ predominant structural details. The resulting data were analyzed using VMD and the figures for reporting were developed using Biovia Discovery Studio Visualizer [27]. The plots of processed data were generated using origin.

## 3. Results

The RBD of 6M0J.PDB contains residues 333 to 526. The RBD mutations on 6M0J are displayed in Fig.1 A-E. Fig. 1A includes the 3 mutations within the Delta/ B.1.617.2 strain selected for this study. Fig. 1B includes the 15 selected mutations within the Omicron/B.1.1.529 strain. Fig. 1C displays the 18 selected mutations within the Omicron BA.2 subvariant. Fig.1D and 1E represents the RBD mutations within the subvariants BA.4 or BA.5 and BA.2.12.1 respectively. Fig.1F-H depicts the S1 mutations in Delta, Omicron variants and Omicron subvariant BA.2. Table S1 and S2 list the descriptive features of these mutations. The mutations within mu variant, subvariants BA.2.75, BF.7 along with a potential subvariant with two mutations are also tabulated there. The RBD mutations of other previously circulating variants have been described in our previous report [16].

**Figure 1.**
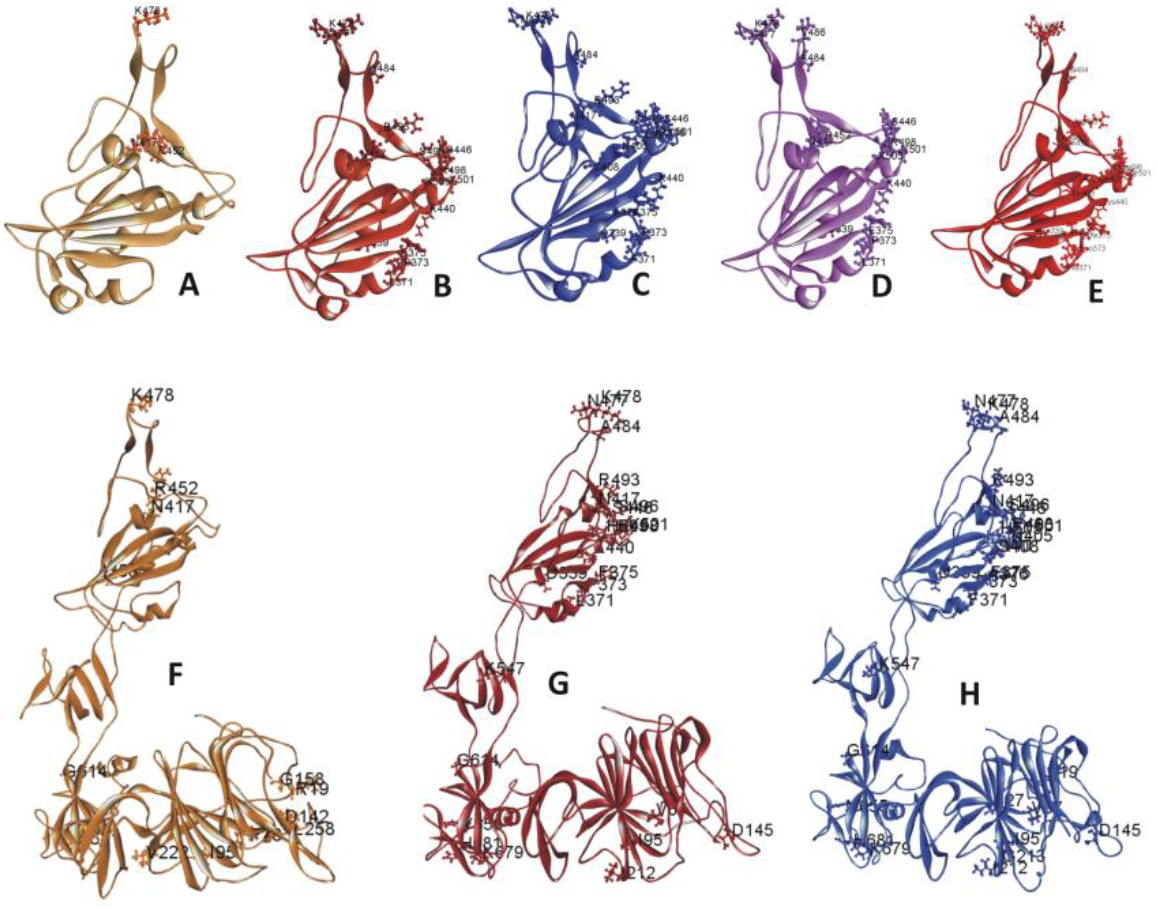
A-E Ribbon diagram of SARS-CoV-2 variants and subvariants with selected RBD mutations. These variants are based on 6M0J structure. Displayed are the RBD of A. Delta variant B. Omicron variant C. Omicron BA.2 subvariant. D. Omicron BA.4/BA.5 subvariant and E. Omicron BA.2.12.1 subvariant with selected RBD mutations F-H. Ribbon diagram of SARS-CoV-2 variants and subvariant with selected S1 mutations. The mutant variants and subvariant structures are based on model S1. Displayed are the S1 of F. Delta variant G. Omicron variant and the H. subvariant BA.2 with selected S1 mutations.

Fig. 2A-B illustrates the traditional/average of all atom RMSD plots of wt and different SARS-CoV-2 variants and subvariant with selected RBD mutations; wt SARS-CoV-2 has also been included here for comparison. Fig. 2C-D presents the traditional and averaged RMSD plots of RBD mutations within different variants. It is evident from RMSD plots (Fig. 2) that Delta RBD and Delta mutations are the most stable form among all species studied here.

**Figure 2.**
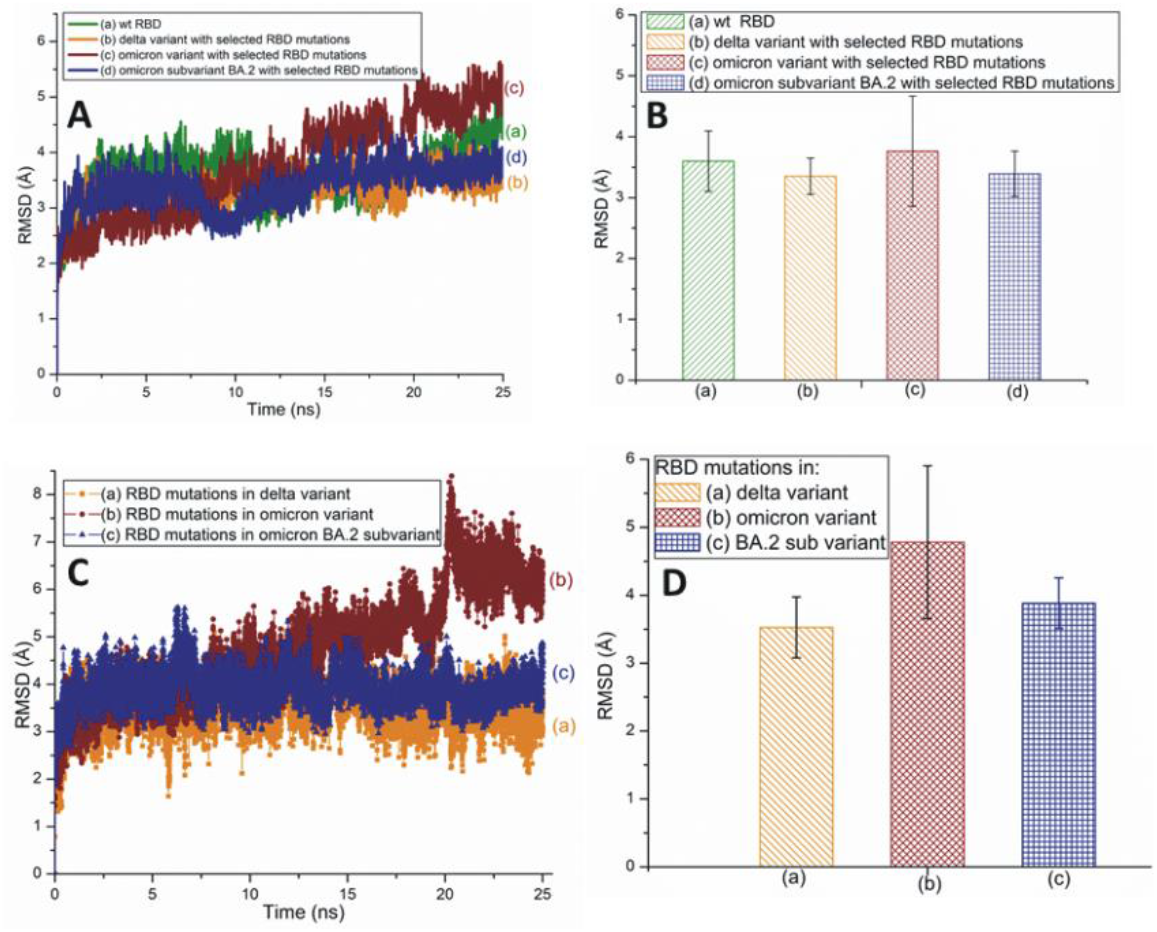
A. The typical all atom RMSD plots of wt and different SARS-CoV-2 variants and subvariant with selected RBD mutations. These variants and subvariant are based on 6M0J structure. B The average RMSD values of wt RBD, Delta, Omicron and BA.2 subvariant with selected RBD mutations. These values are extracted from Fig. 2A. The average RMSD value was taken from a 25ns simulation time. C. The typical RMSD plots of RBD mutations within different SARS-CoV-2 variants and subvariant. D. The average RMSD values of SARS-CoV-2 mutations within deferent variants and subvariant. These values are extracted from Fig. 2C. In all cases the all atom RMSD values are considered.

Fig. 3A illustrates traditional all atom RMSD plots of the SARS-CoV-2 Omicron subvariants, BA.4/BA.5 and BA.2.12.1 with selected RBD mutations. The average RMSD values of these subvariants are displayed in Fig. 3B. The all atom RMSD values of RBD mutations within these two subvariants are plotted in Fig. 3C-3D. A comparison of the RMSD values of BA.4/BA.5 and a possible subvariant with two mutations L452R and F486V are plotted in SM Fig. S1. The RBD RMSD graphs of Mu variant and BA.2.75 subvariant are assembled in Fig. S2. The RMSD plot of BF.7 RBD with selected mutations is plotted in Fig. S3. The RMSD of mutations within BF.7 is also plotted there. The RMSD of mutation R346T, observed in BF.7 is plotted in Fig. S3B [6,28]. The RMSD plots of circulating VOI and VUMs as listed by WHO are displayed in SM Fig. S4. The RBD root-mean-square fluctuation (RMSF) plots of variants and subvariants are plotted in Fig. S5.

**Figure 3.**
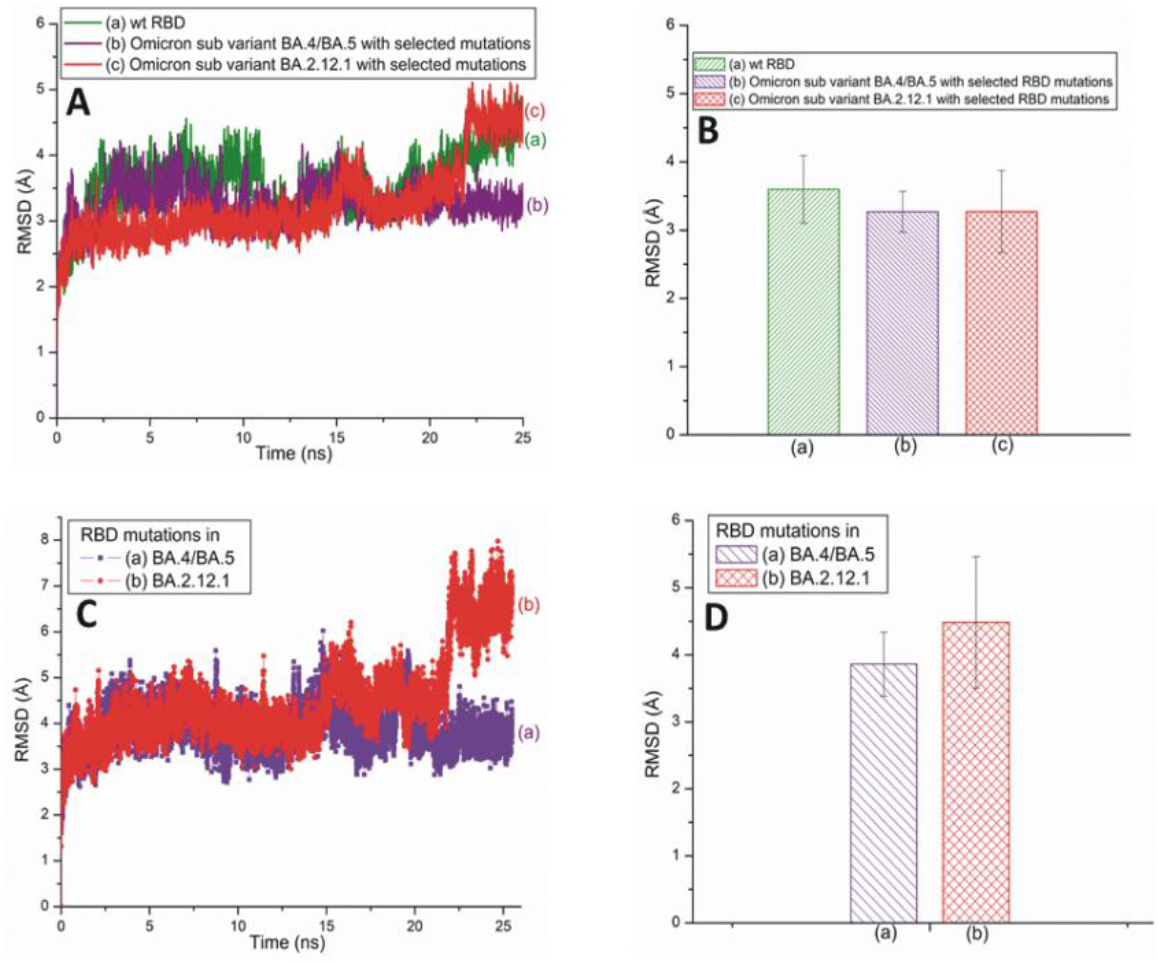
A. The typical all atom RMSD plots of wt and recently emerged SARS-CoV-2 Omicron subvariants with selected RBD mutations. These subvariants are based on 6M0J structure. B The average RMSD values of wt RBD, subvariants BA.4/BA.5 and BA.2.12.1 with selected RBD mutations. The values were taken from the 25 ns simulation time. These values were extracted from Fig. 3A. C. The typical all atom RMSD plots of RBD mutations within subvariants BA.4/BA.5 and BA.2.12.1. D. The average RMSD values for SARS-CoV-2 mutations within deferent subvariants. The average values were extracted from Fig. 3C.

Fig. 4A-B represents the traditional and averaged all atom RMSD graphs of the wt and the mutant variants/subvariant of SARS-CoV-2 S1 with selected mutations. Fig 4C-D shows the traditional and averaged RMSD values of the S1 mutations within selected variants/subvariant. The RMSD plot of Mu S1 is plotted in Fig, S6. The RMSF of wt and mutant S1 is plotted in Fig. S7A. A zoomed-in view of plot S1 is displayed in Fig. S7B.

**Figure 4.**
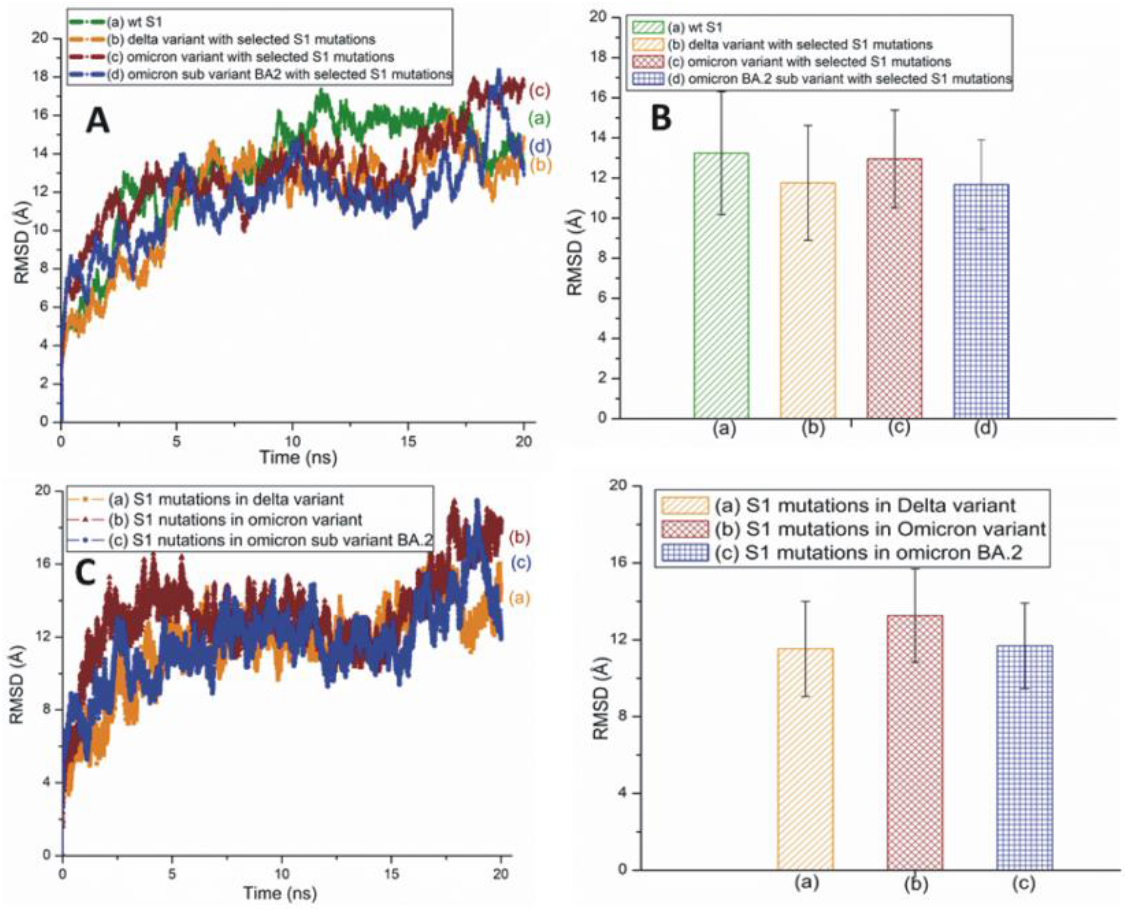
A. The typical all atom RMSD plots of wt and different SARS-CoV-2 variants and subvariant with selected S1 mutations. These variants and subvariant are based on model structure of SARS-CoV-2 S1. B The average RMSD values of wt S1, the Delta and Omicron variants and Omicron subvariant BA.2 with selected S1 mutations. The average RMSD value was taken from a 20ns simulation time. These values were extracted from Fig. 4A. C. The typical RMSD plots of S1 mutations within different SARS-CoV-2 variants and subvariant. D. The average RMSD values of SARS-CoV-2 S1 mutations within deferent variants. The average values were extracted from Fig. 4C.

Results of time based secondary structure analyses of different RBDs are displayed in Fig. 5. Time based secondary structures of few more variant and subvariants RBDs are displayed in Figs. S8-S10. Fig. 6 illustrates the secondary structure changes of SARS-CoV-2 S1 of Delta, and Omicron variants and BA.2 subvariant with selected S1 mutations. The secondary structure changes of SARS-CoV-2 S1 within Mu variant is plotted in Fig. S11.

**Figure 5.**
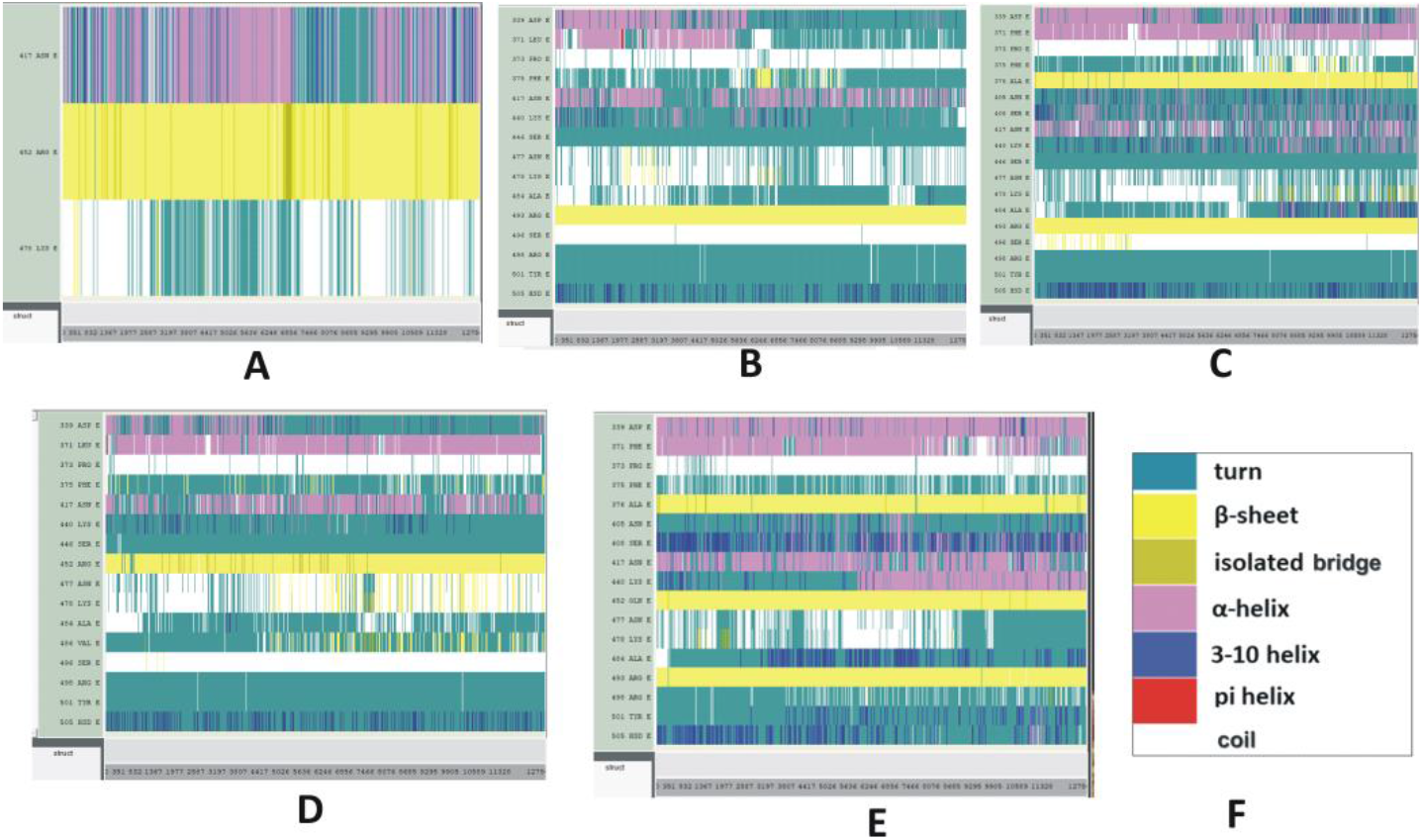
Time based secondary structure changes of SARS-CoV-2 mutant variants and subvariants. The vertical axis represents the mutated residues and the horizontal axis represents the simulation trajectory. A-E. the SARS-CoV-2 RBD of (A) Delta, (B) Omicron variants and Omicron subvariants (C) BA.2 (D) BA.4/BA.5 (E) BA.2.12.1 with selected RBD mutations. These secondary structural changes were recorded for 25ns. F. Color code explanation of proteins’ secondary structures. These are the default color codes generated by VMD GUI.

**Figure 6.**
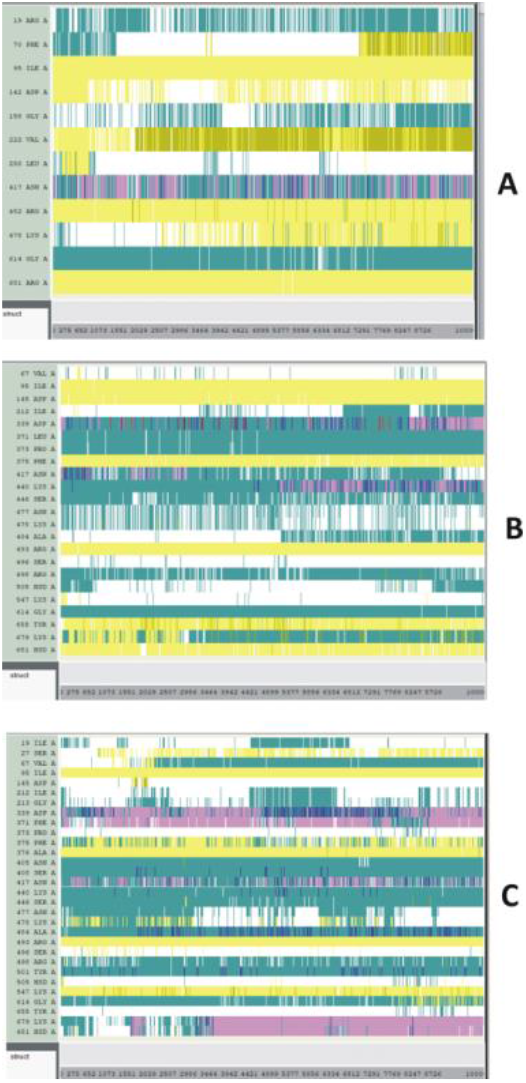
The Secondary structure changes of SARS-CoV-2 S1. A. Delta and B. Omicron variants and C. Omicron subvariant BA.2 with selected S1 mutations. The structural changes were recorded for 20ns.

The average solvent accessible surface area (SASA) values for each of the RBD mutations within the SARS-CoV-2 variants and subvariants are plotted in Fig.7.

The SASA values are calculated using the VMD timeline window. These results are average SASA values obtained from the simulation time scale. The SASA values of S1 mutations are displayed in Fig 8A-C, where the findings of the corresponding RBDs are detected again.

**Figure 7.**
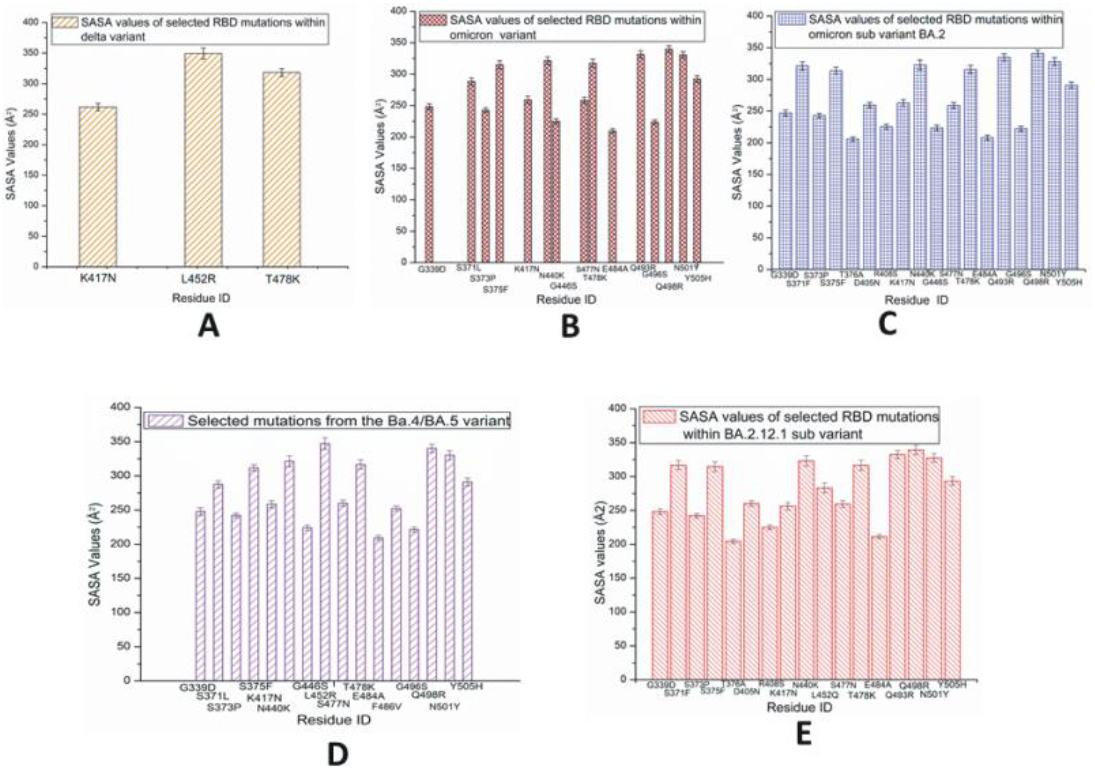
Average SASA values of SARS-CoV-2 RBD mutations observed predominantly in variants A. Delta; B. Omicron and subvariants C. BA.2; D. BA.4/BA.5; and E. BA.2.12.1.

**Figure 8.**
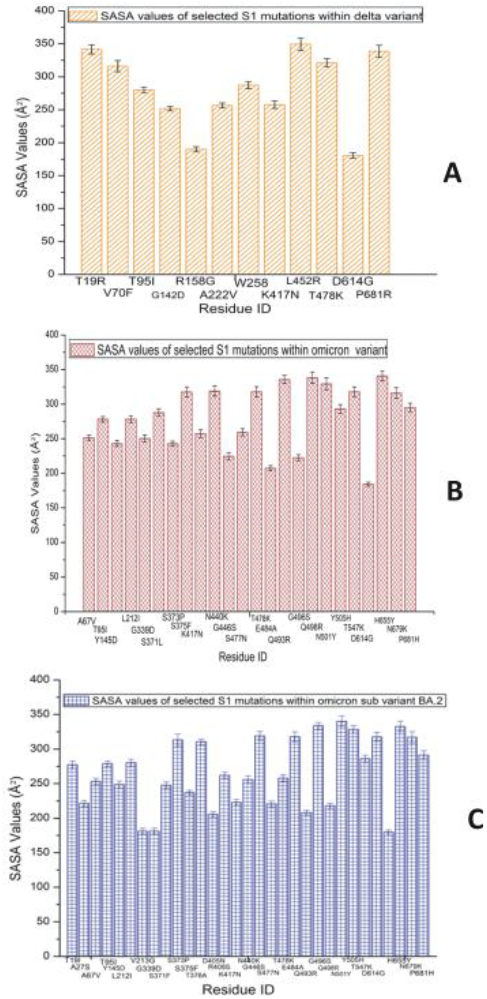
Average SASA values of SARS-CoV-2 S1 mutations observed predominantly in A. Delta variant; B. Omicron variant; and C. Omicron subvariant BA.2.

## Discussion

Fig.1 depicts the secondary structure of RBD and S1 of various SARS-CoV-2 variants and subvariants. Since most of the mutations in BA.4/BA.5 and BA2.12.1 are concentrated and hence observed in their RBDs, we have specifically examined the RBDs of these three subvariants in the present report.

The Delta variant is more stable than Omicron and even stable compare to its wt version (Fig. 2A). While the Omicron variant is most unstable, the RBD of BA.2 is also quite stable. The average root mean square deviation (RMSD) of the BA.2 subvariant is slightly higher (3.4Å) than that of the Delta variant (3.35Å) as displayed in Fig. 2B. Fig. 2B also describes the average all atom RMSD results within the wt (3.6Å) and the Omicron (3.76 Å) species. The Delta variant has the lowest average RMSD value while Omicron has the highest. This figure also indicates certain findings from Fig. 2A, where the Omicron variant shows larger variations. The results in Figs 2C-2D suggest that the mutations within the Delta variant (3.52Å) are most stable and those of Omicron are most unstable (4.78Å). It is possible that numerous RBD mutations within the Omicron make the latter unstable; this is not the case for the RBD of Omicron subvariant BA.2, where the presence of RBD mutations make this protein rather stable (3.88Å). While the earlier version of the BA.2 has been examined in this work, the recently updated information from the CDC should be noted in this context, according to which, the BA.2 belongs to Omicron [2].

From Fig.3A-B it is evident that RBD RMSD of subvariant BA.4/BA.5 (RMSD value: 3.26Å) is almost equal to that of BA.2.12.1 (RMSD value:3.27Å). However the RMSD values of the mutations observed within these two subvariants demonstrated that the BA.4/BA.5 mutations have a lower RMSD value of 3.86Å compare to 4.48Å of BA2.12.1. Accordingly, the mutations within BA.4/BA.5 are more stable than those existing within BA.2.12.1. This also implies that the mutations within BA.4/BA.5 are unstable compare to its entire RBD (Figs. 3 and S1). Therefore, though the BA.4/BA.5 RBD is more stable than Delta RBD overall, the RBD mutations within Delta (RMSD value: 3.52Å) are more stable than those of BA.4/BA.5 (3.86Å) and Omicron (4.78Å) variants, as well as those of BA.2 (3.88Å) and BA.2.12.1 (4.48Å) subvariants (Figs 2C-D and 3C-D). Although the mutations in BF.7 exhibit higher RMSD value, it is evident from these figures that mutation R346T within BF7 subvariant is reasonably stable (Fig. S3). Fig. S4 suggests that XBF subvariant has the lowest RMSD value (3.5) among all the listed VOI and VUMs; it is possible that the vaccine and booster doses are able to control XBF subvariant of SARS-CoV-2 so far. The RMSF plot (Fig. S5) mimics the RMSD data, where the delta RBD is most stable. Additionally, the RMSF data indicate that omicron subvariants BA.4/BA.5 and BA2 have stable RBDs. The RMSF plot for BF.7 RBD is rather stable except in the cases of residues 474-486 (Fig. S5B). Among these residues, 482-483 display highest instability. It is possible that the presence of several mutated residues (477, 478, 484 and 486) near 482-483, and a lack of proper folding lead to a structurally disordered region [29]. Thus, in general, the overall trend for this particular S protein region is unstable and variable, apparently, due to the presence of local loops and turns (Fig. S5B). As mutations serve as markers of genetic variations, it is difficult to quantify their stability; however, mutational effects on the proteins’ general stability can be predicted. A recently published article has demonstrated this point, where the S protein mutations within Omicron variant have been found to play roles in allosterically regulating the stability, conformational flexibility and structural adaptability of the protein [30].

Based on Fig.4, we can assume that S1 of Delta and BA.2 are the stable form of this virus among these variants and subvariant. The overall all atom RMSD values of the S1 are higher than the RBD RMSDs due to the presence of many turns and coils in this segment. From the S1 RMSF plots (Fig. S7) it is clear that Omicron S1 residues have high RMSF values compare to the delta and BA.2 S1. This is particularly prominent in the RBDs of the S1 system displayed in Fig. S7B, a zoomed-in view of Fig. S7A.

Within the Delta variant, the secondary structure of K417N and L452R mutations are stable compared to T478K (Fig. 5A). In the Omicron variant most of the mutations are unstable including G339D, S371L, S375F, N440K, S477N and T478K, which, in combination, make the Omicron RBD measurably unstable (Fig. 5B). In BA.2 subvariant R408S, T478K and E484A residues seems very stable whereas residues G339D and G496S appear unstable (Fig.5C). These results, suggest an overall high stability of the Delta RBD. Based on its RBD, the subvariant BA.2 is also a stable protein and Omicron RBD is notably unstable among these three RBDs.

Most of the mutations identified in BA.4/BA.5, displayed in Fig.5D, are quite stable. Fig. 5E represents the secondary structure changes of the RBD mutations within BA.2.12.1 subvariant. While mutation R408S appears stable with time, according to the present results, mutations S371F, K417N, T478K and Y505H do not seem to follow the same the same trend. It should be noted in this context that though mutation S477N and T478K are slightly stable in the subvariant BA.4/BA.5, these two mutations are unstable within Omicron variant and subvariant BF.7. However mutation R346T within BF.7 does not indicate any major conformational changes in its secondary structure during the course of the simulation process (Fig. S9B).

The Delta S1 appears quite stable (Fig.6A). No major changes in the stability or fluctuations within the secondary structures are observed for this variant. V70F and T478K residues also seem quite stable during the simulation timescale as the turns and coils of these species are converted into beta sheets. In the case of Omicron S1, though residues G339D and N440K seem stable with time, fluctuations are observed in the mutated residues A67V, L212I, K417N and T547K (Fig. 6B). For the BA.2 subvariant (Fig.6C), residues A67V and T478K seem unstable as their beta sheets are converted into turns and coils. On the other hand, residues S375F and residues 479-81 are converted to alpha helices and beta sheets from their turns/coils phases, respectively. The stability analyses of S1 secondary structure mimics those of the RBD secondary structures, and suggests that Delta is the most stable variant.

In Fig. 7A, the highest SASA values are observed in L452R within the RBD of Delta variant, which suggests that this residue has a comparably large solvent accessible surface. It should be noted in this context that the severity of certain disease largely depends on the buried or exposed nature of the underlying mutated residues [31]. The SASA values may increase due to partial unfolding caused by residue mutation. Among the three species we have studied here, the L452R mutation is only observed in the Delta and few subvariants. Since mutated residue R is hydrophilic and surface exposed, the structural rearrangements or conformational deviations may possible in this case. Published report suggests that L452R within the Delta variant strongly interacts with ACE2 [32]. T478K within Delta variant also shows considerably high SASA value. As positively charged K^+^ impact the electrostatic surface area, this residue may manifest stronger interactions with the receptor [33]. In Delta RBD, only one residue shows considerably lower SASA values indicating that this buried residue K417N is quite stable in nature. However, if surface exposed residues on the RBD form stable interactions with the ACE2 receptor as in the case of Delta, where the mutant L452R strongly bind to the ACE2 receptor making Delta RBD quite stable. Thus it could be possible that the L452R and T478K within the Delta variant may have a stronger stabilizing effect [16,34]. Although T478K was not present in any previously identified variants, this mutation is observed in all the variants and subvariant we studied in this report. It is possible that the greater mutational fitness of the Delta variant may be due to L452R or a combination of a number of residue mutations. In the case of Omicron RBD, as displayed in Fig. 7B, despite the presence of fewer buried residues with lower SASA values, majority of the mutated residues have greater SASA values; this suggests the possibility of making this protein quite unstable [35]. The Omicron variant contains more mutations than Delta and some of them are at the ligand-receptor interfacial region, while the stronger binding interactions between L452R and ACE2 receptor make Delta variant somewhat unique. Hydrogen bond formation between residue Q493 of Delta with ACE2 contributes to its stronger structure [32]. Furthermore, it is possible that most of the mutations present in the Omicron variant are counteracted by neutralizing antibodies, while at the same time, L452R in Delta is considered as antibody neutralizing escape mutation. The Omicron and its subvariant BA.4/BA.5 have a lower hydrophobic potential compare to most of the other previously identified variants. Omicron’s RBD including the Omicron subvariant BA.4/BA.5 consists of a positive electrostatic surface, while the ACE2 receptor contains many negatively charged residues; therefore, strong binding due to coulombic interactions at the protein-protein interface region is possible [36]. Moreover, since these positively charged RBD mutations exist within the antibody neutralization epitope regions, they could serve to facilitate immune escape. The charged and hydrophobic surface map of BA.4/BA.5 is displayed in Fig. S12. In the case of BA.2, as displayed in Fig.7C the slightly higher numbers of buried residues (with lower SASA values) possibly make this subvariant more stable than original Omicron. Usually, the buried residues present within the core of the protein are the most stable part of a protein. The surface exposed residues with greater SASA values are likely to be comparatively less stable unless the surface exposed residues form stronger receptor interactions. Fig. 7D represents the SASA values of RBD mutations within BA.4/BA.5. Here the additional signature mutation L452R demonstrates higher SASA values compared to the other signature mutation, F486V. As discussed before, L452R plays a key role in receptor residue interaction. Among the other strains studied in this report, mutation L452R is only present in the Delta and BA.2 variant. In BA.2.12.1, the 452 mutation is denoted as L452Q. It is possible that the BA.5 subvariant has become a dominant strain due to the antibody neutralization escape nature of L452R. Considering that the RBDs of BA.4 and BA5 contain mutually similar mutations, certain structural/mutational differences within the S protein may exist that make BA.5 more transmissive than BA.4. Fig. S13 represents the SASA values of a possible subvariant with two mutations L452R and F486V. These two mutations are observed in BA.4/BA.5 in addition to the mutations that are observed in Omicron BA.2. The SASA values of RBD mutations in BA.2.12.1 are displayed in Fig. 7E. Although some relatively low SASA values are observed, majority of the BA.2.12.1 mutations are surface exposed and contain greater SASA values. Overall, BA.2 subvariant has more buried residues than those of BA.212.1.

Within the S1 (Fig. 8A) The Delta has a number of buried residues, but here the surface exposed residues have strong interactions with ACE2. Despite the many buried residues of Omicron S1 (Fig. 8B), the numbers of surface exposed residues are greater than the buried residues in this case. S1 of subvariant BA.2 (Fig. 8C) has more buried residues than those of the S1 of Omicron variant. It is likely that most of the surface exposed residues in the Omicron variant are loosely attached to the ACE2 receptor. Currently there are no readily available experimental results for the BA.2 subvariant. This subvariant may have antibody neutralizing escape mutants and thus, may emerge as a strong form of the virus [37,38]. Nonetheless, according to the RMSD data and the time-based secondary structure analyses of S1 data presented here, the Delta and subvariant BA.2 are the stable forms among the three forms we have studied. For the RBD structure, BA.4/BA.5 is also quiet stable along with them.

Omicron subvariant BA.2 has been known as “stealth Omicron” as this subvariant was initially difficult to identify from the Delta variant using traditional testing procedures. A combination of subvariants BA.1 and BA.2 has been also identified. This hybrid or recombinant variant is also known as subvariant XE [39]. BA.5 sublineage BF.7, has been reported in some countries. Based on the limited information currently available it could be predicted that BF.7 has higher spreadibility with short incubation time and greater antibody neutralization escape efficacy possibly due to stable RBD mutation R346T in addition to the BA.5 mutations [40] Glycosylation sites are important as they play critical roles in immune evasion. Since BF.7 has mutation R346T close to N331 and N343 (two N-linked glycosylation sites of S1 RBD), that could possibly be linked to higher immune escape [36,41]. It could be noted that some of the circulating VOI and VUMs also have mutations close to the N-linked glycosylation sites within their S1 RBD (Table S3).

## 4. Conclusions

The simulation data reported here indicate that Delta is a stable form of COVID 19. However, among the forms selected for the present study, BA.4/BA.5 and BA.2 RBD also appears quiet stable. Additionally, some of the mutations discussed in this report play crucial role in formation of stable variants/subvariants including XBB, a circulating VUM. Recently the mRNA bivalent boosters have been released in US that contain mRNA of original wt strain as well as BA.4/BA.5 subvariants and thus are more effective against aggressive infections. However as RNA virus is highly adaptable and environmentally fit, it is possible that newer versions of variants/subvariants with new functional epitopes may emerge with possible breakthrough infections. We expect that the detailed structural data discussed in this paper will be useful to identify drug targets and neutralizing antibodies for emerging SARS-CoV-2 variants or subvariants. We also hope that by contributing to the literature on SARS-CoV-2, the present manuscript will serve to help the ongoing efforts to overcome the COVID 19 pandemic in some ways.

## Supplementary Materials

Figs S1-S13; Table S1-S3

## Conflict of interest statement

The author declares no conflict of interest.

## Funding

The authors declare that no funds, grants, or other support were received duringthe preparation of this manuscript

## Acknowledgments

The author acknowledges utilization of the following simulation and visualization software packages: 1) NAMD and 2) VMD: NAMD and VMD, developed by the Theoretical and Computational Biophysics Group in the Beckman Institute for Advanced Science and Technology at the University of Illinois, Urbana-Champaign. 3) Discovery Studio Visualizer: Dassault Systèmes BIOVIA, Discovery Studio Modeling Environment, San Diego, CA: Dassault Systèmes (2015).

## Notes

### Competing Interest Statement

The authors have declared no competing interest.

### Summary of Updates

Currently circulating variants of interest and variants under monitoring including XBB.1.5, BQ.1/BQ.1.1, CH.1.1, XBB and XBF are briefly explored in this version.

## References

1. WHO, Weekly epidemiological update (2021). https://www.who.int/publications/m/item/weekly-epidemiological-update23-february-2021.Accessed 25 February 2021

2. Centers for Disease Control and Prevention, SARS-CoV-2 Variant Classifications and Definitions. https://www.cdc.gov/coronavirus/2019-ncov/variants/variant-info.html#Interest. Accessed 22 June 2021, 26 July 2021, 7 December 2021, 26 April 2022, June, 2022

3. Science Brief: Omicron (B.1.1.529) Variant. https://www.cdc.gov/coronavirus/2019-ncov/science/science-briefs/scientific-brief-omicron-variant.html. Accessed 18 February 2022

4. Variant: 21L (Omicron) also known as BA.2. https://covariants.org/variants/21L.Omicron. Accessed 22 March 2022

5. SARS-CoV-2 variants of concern as of 9 June 2022. https://www.ecdc.europa.eu/en/covid-19/variants-concern Accessed 17 June 2022, 12 July 2022

6. Tracking SARS-CoV-2 variants. https://www.who.int/en/activities/tracking-SARS-CoV-2-variants/. Accessed 14 June 2022, 18 January 2023

7. Haspel, N.; Jang, H.; Nussinov, R. (2021) Active and Inactive Cdc42 Differ in Their Insert Region Conformational Dynamics. Biophys. J. 120: 306–318

8. Huang, C.J.; Schild, L.; Moczydlowski, E.G. (2012) Use-dependent block of the voltage-gated Na(+) channel by tetrodotoxin and saxitoxin: effect of pore mutations that change ionic selectivity. J Gen Physiol 140: 435–454

9. Lampe, A.T.; Puniya, B.L.; Pannier, A.K.; Helikar, T.; Brown, D.M. (2020) Combined TLR4 and TLR9 agonists induce distinct phenotypic changes in innate immunity in vitro and in vivo. Cell. Immunol. 355: 104149

10. Kaya, T.; Swamy, N.; Persons, K.S.; Ray, S.; Mohr, S.C.; Ray, R. (2009) Covalent labeling of nuclear vitamin D receptor with affinity labeling reagents containing a cross-linking probe at three different positions of the parent ligand: structural and biochemical implications. Bioorg Chem 37: 57–63

11. Farooq, M.; Khan, A.W.; Ahmad, B.; Kim, M.S.; Choi, S. (2022) Therapeutic Targeting of Innate Immune Receptors Against SARS-CoV-2 Infection. Front Pharmacol 13: 915565

12. Roy, U. (2016) Structural Characterizations of the Fas Receptor and the Fas-Associated Protein with Death Domain Interactions. The Protein J. 35: 51–60

13. Roy, U. (2019) 3D Modeling of Tumor Necrosis Factor Receptor and Tumor Necrosis Factor-bound Receptor Systems. Mol. Inform. 38: e1800011

14. Roy, U. (2020) Structural and molecular analyses of functional epitopes and escape mutants in Japanese encephalitis virus envelope protein domain III. Immunol. Res. 68: 81–89

15. Roy, U. (2021) Role of N501Y mutation in SARS-CoV-2 spike proteinstructure. Preprints, 2021060238

16. Roy, U. (2021) Comparative Structural Analyses of Selected Spike Protein-RBD Mutations in SARS-CoV-2 Lineages. Immunol. Res. 70: 143–151

17. Fatouros, P.R.; Roy, U.; Sur, S. (2022) Modeling Substrate Coordination to Zn-Bound Angiotensin Converting Enzyme 2. Int. J. Pept. Res. Ther. 28: 65

18. Wang, Q.; Guo, Y.; Iketani, S.; Nair, M.S.; Li, Z.; Mohri, H.; Wang, M.; Yu, J.; Bowen, A.D.; Chang, J.Y.; et al. (2022) Antibody evasion by SARS-CoV-2 Omicron subvariants BA.2.12.1, BA.4 and BA.5. Nature 608: 603–608

19. Lan, J.; Ge, J.; Yu, J.; Shan, S.; Zhou, H.; Fan, S.; Zhang, Q.; Shi, X.; Wang, Q.; Zhang, L.; et al. (2020) Structure of the SARS-CoV-2 spike receptor-binding domain bound to the ACE2 receptor. Nature 581: 215–220

20. Yang, J.; Zhang, Y. (2015) I-TASSER server: new development for protein structure and function predictions. Nucleic Acids Res. 43: W174–181

21. Walls, A.C.; Park, Y.J.; Tortorici, M.A.; Wall, A.; McGuire, A.T.; Veesler, D. (2020) Structure, Function, and Antigenicity of the SARS-CoV-2 Spike Glycoprotein. Cell 181: 281–292.e286

22. Xiong, X.; Qu, K.; Ciazynska, K.A.; Hosmillo, M.; Carter, A.P.; Ebrahimi, S.; Ke, Z.; Scheres, S.H.W.; Bergamaschi, L.; Grice, G.L.; et al. (2020) A thermostable, closed SARS-CoV-2 spike protein trimer. Nat. Struct. Mol. Biol. 27: 934–941

23. Humphrey, W.; Dalke, A.; Schulten, K. (1996) VMD: Visual molecular dynamics. J. Mol. Graph. 14: 33–38

24. Phillips, J.C.; Braun, R.; Wang, W.; Gumbart, J.; Tajkhorshid, E.; Villa, E.; Chipot, C.; Skeel, R.D.; Kale, L.; Schulten, K. (2005) Scalable molecular dynamics with NAMD. J. Comput. Chem 26: 1781– 1802

25. Ribeiro, J.V.; Bernardi, R.C.; Rudack, T.; Stone, J.E.; Phillips, J.C.; Freddolino, P.L.; Schulten, K. (2016) QwikMD - Integrative Molecular Dynamics Toolkit for Novices and Experts. Sci. Rep. 6: 26536

26. Tanner, D.E.; Phillips, J.C.; Schulten, K. (2012) GPU/CPU Algorithm for Generalized Born/Solvent-Accessible Surface Area Implicit Solvent Calculations. J. Chem. Theory Comput. 8: 2521–2530

27. Dassault Systèmes BIOVIA Discovery Studio Modeling Environment, San Diego, CA: Dassault Systèmes (2015).

28. SARS-CoV-2 variants of concern and variants under investigation in England, Technical briefing 46. https://assets.publishing.service.gov.uk/government/uploads/system/uploads/attachment_data/file/1115070/Technical-Briefing-46-7October2022.pdf. Accessed 22 December 2022

29. Quaglia, F.; Salladini, E.; Carraro, M.; Minervini, G.; Tosatto, S.C.E.; Le Mercier, P. (2022) SARS-CoV-2 variants preferentially emerge at intrinsically disordered protein sites helping immune evasion. The FEBS Journal 289: 4240–4250

30. Verkhivker, G.M.; Agajanian, S.; Kassab, R.; Krishnan, K. (2022) Frustration-driven allosteric regulation and signal transmission in the SARS-CoV-2 spike omicron trimer structures: a crosstalk of the omicron mutation sites allosterically regulates tradeoffs of protein stability and conformational adaptability. Physical Chemistry Chemical Physics 24: 17723–17743

31. Savojardo, C.; Manfredi, M.; Martelli, P.L.; Casadio, R. (2020) Solvent Accessibility of Residues Undergoing Pathogenic Variations in Humans: From Protein Structures to Protein Sequences. Front Mol. Biosci. 7: 626363

32. Socher, E.; Heger, L.; Paulsen, F.; Zunke, F.; Arnold, P. (2022) Molecular dynamics simulations of the delta and omicron SARS-CoV-2 spike - ACE2 complexes reveal distinct changes between both variants. Comput. Struct. Biotechnol. J. 20: 1168–1176

33. Di Giacomo, S.; Mercatelli, D.; Rakhimov, A.; Giorgi, F.M. (2021) Preliminary report on severe acute respiratory syndrome coronavirus 2 (SARS-CoV-2) Spike mutation T478K. J. Med. Virol. 93: 5638–5643

34. Goher, S.S.; Ali, F.; Amin, M. (2021) The Delta Variant Mutations in the Receptor Binding Domain of SARS-CoV-2 Show Enhanced Electrostatic Interactions with the ACE2. Med. Drug Discov.: 100114

35. Zeng, C.; Evans, J.P.; Qu, P.; Faraone, J.; Zheng, Y.-M.; Carlin, C.; Bednash, J.S.; Zhou, T.; Lozanski, G.; Mallampalli, R.; et al. (2021) Neutralization and Stability of SARS-CoV-2 Omicron Variant. bioRxiv: 2021.2012.2016.472934

36. López-Cortés, G.I.; Palacios-Pérez, M.; Veledíaz, H.F.; Hernández-Aguilar, M.; López-Hernández, G.R.; Zamudio, G.S.; José, M.V. (2022) The Spike Protein of SARS-CoV-2 Is Adapting Because of Selective Pressures. Vaccines (Basel) 10:

37. Chen, J.; Wei, G.-W. (2022) Omicron BA.2 (B.1.1.529.2): high potential to becoming the next dominating variant. ArXiv: arXiv:2202.05031v05031

38. Kannan, S.R.; Spratt, A.N.; Sharma, K.; Sönnerborg, A.; Apparsundaram, S.; Lorson, C.; Byrareddy, S.N.; Singh, K. (2022) Complex Mutation Pattern of Omicron BA. 2: Evading Antibodies without Losing Receptor Interactions. Int. J. Mol. Sci. 23: 5534

39. COVID-19 subvariant XE: What to know. https://www.foxnews.com/health/covid-19-subvariant-xe-ba-2-coronavirus-pandemic. Accessed accessed May 10, 2022

40. BF.7: What to know about the Omicron COVID variant. https://www.cbsnews.com/news/bf7-new-omicron-coronavirus-variant-covid/. Accessed 22 December 2022

41. Watanabe, Y.; Allen, J.D.; Wrapp, D.; McLellan, J.S.; Crispin, M. (2020) Site-specific glycan analysis of the SARS-CoV-2 spike. Science 369: 330–333

