## Supplementary material for "Molecular Investigations of Selected Spike Protein Mutations in SARS-CoV-2: Delta and Omicron Variants and Omicron Subvariants": Figs S1-S13; Table S1-S3

*Urmi Roy*[*]

[*]Department of Chemistry & Biomolecular Science

Clarkson University

8 Clarkson Avenue, Potsdam, NY 13699-5820, United States

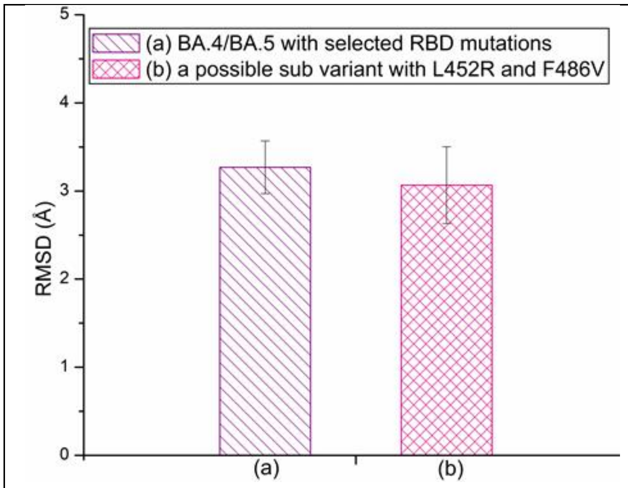


(A)

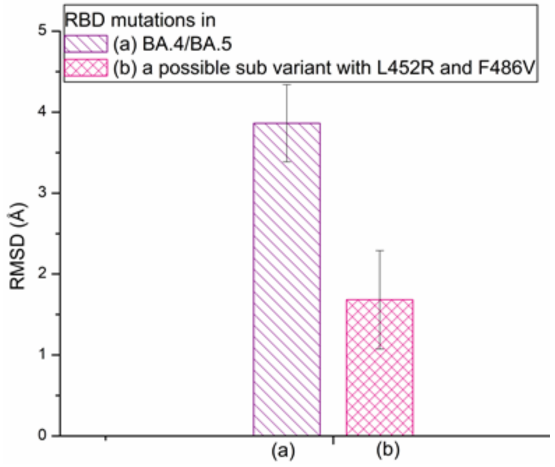


(B)

**Fig. S1.A.** The average RMSD plots for subvariant (a) BA.4/BA.5 and (b) a possible variant with two mutations L452R and F486V. B. The average RMSD plots of mutations within (a) subvariant BA.4/BA.5 and (b) a possible subvariant with two mutations L452R and F486V. These two mutations are observed in the existing BA.4/BA.5 subvariant in addition to the BA.2 mutations.

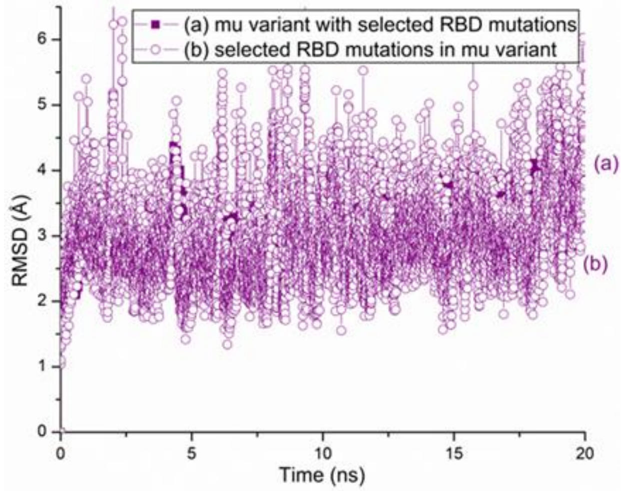


(A)

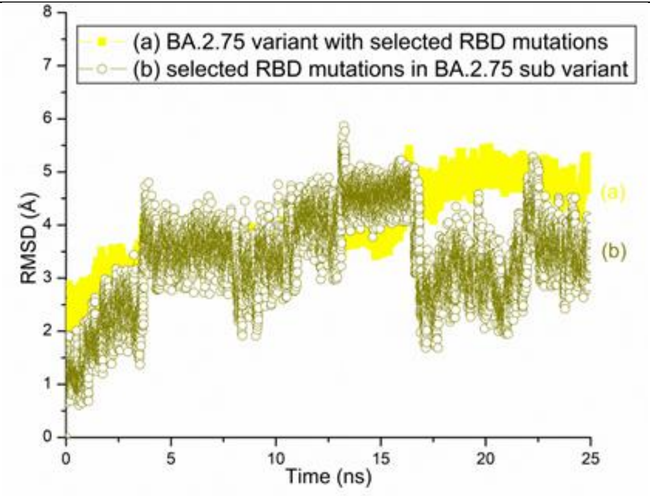


(B)

**Fig. S2.** A. The RMSD plots for the SARS-CoV-2 (a) mu variant RBD and (b) RBD mutations within the mu variant. B. The RMSD plots of (a) BA.2.75 subvariant (b) and RBD mutations in BA.2.75 sub variant.

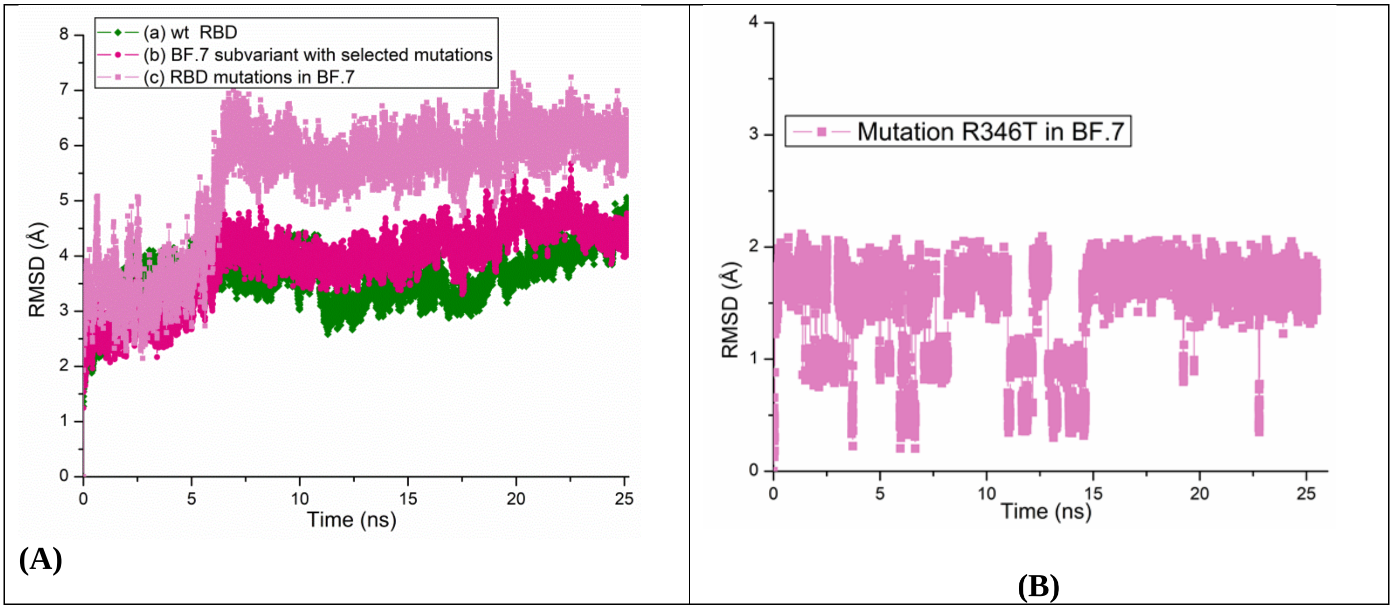


**Fig. S3.** A. The all atom RMSD plots for the SARS-CoV-2 (a) wt protein (b) BF.7 subvariant with RBD mutations and (c) RBD mutations within BF.7 subvariant. B. The RMSD plots of R346T mutation in BF.7 subvariant

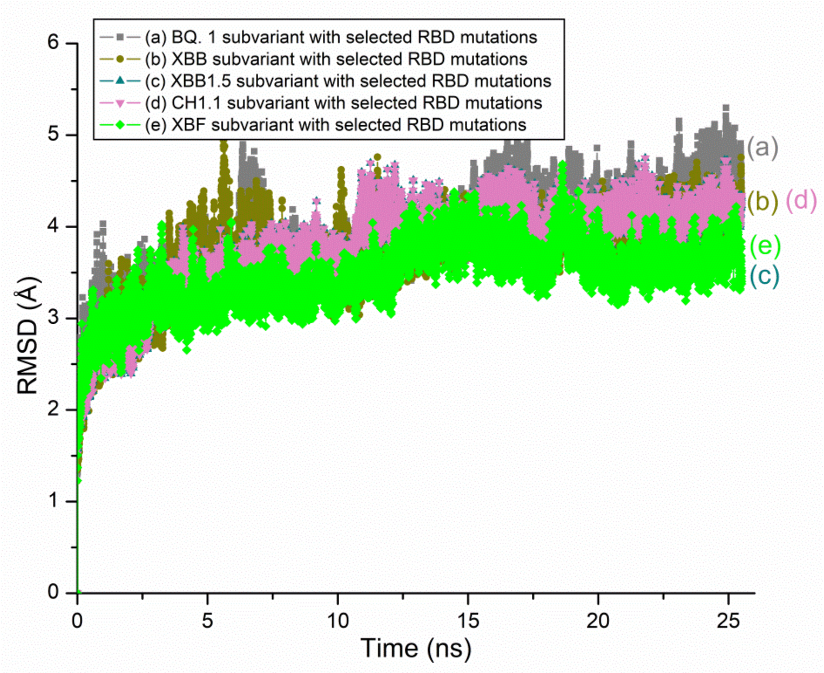


**Fig. S4.** The all atom RMSD plots for the circulating VOI and VUMs of SARS-CoV-2 RBD. In all cases, the variant proteins RBDs are based on 6M0J.PDB.

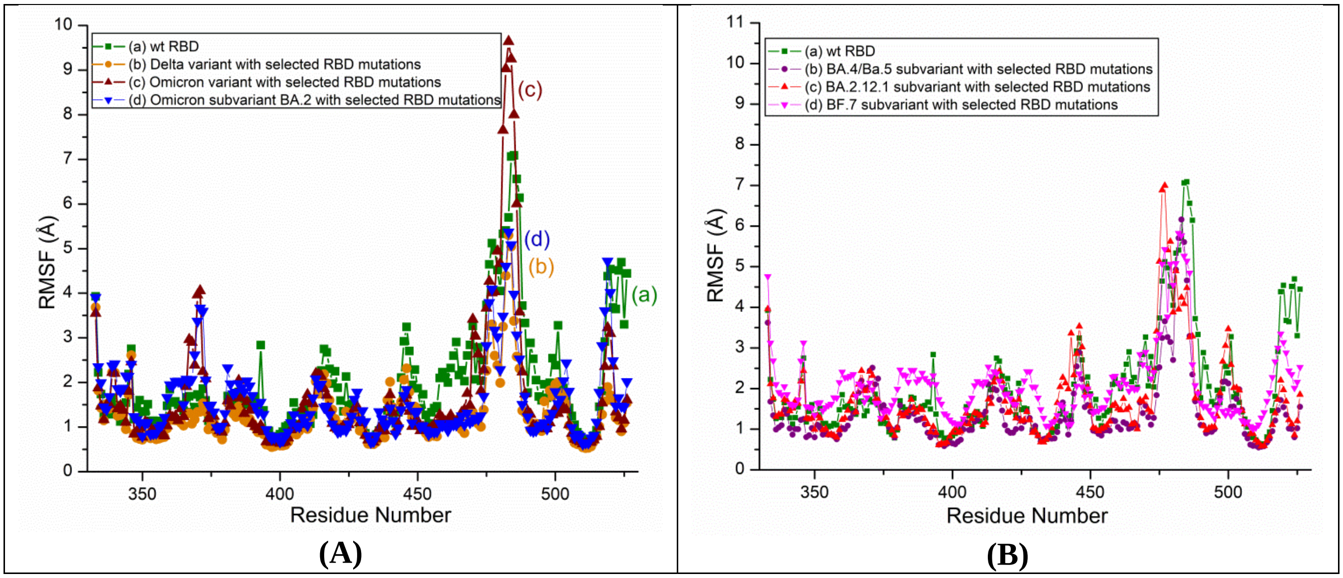


**Fig. S5.** RMSF plots of SARS-CoV-2 variants and subvariants. A.The RMSF plots of wt RBD, Delta RBD, Omicron RBD and BA.2 s RBD with selected mutations. B. The RMSF plots of wt RBD and RBDs of subvariants BA.4/BA.5, BA.2.12.1 and BF.7 with selected mutations. These subvariants are based on 6M0J structure.

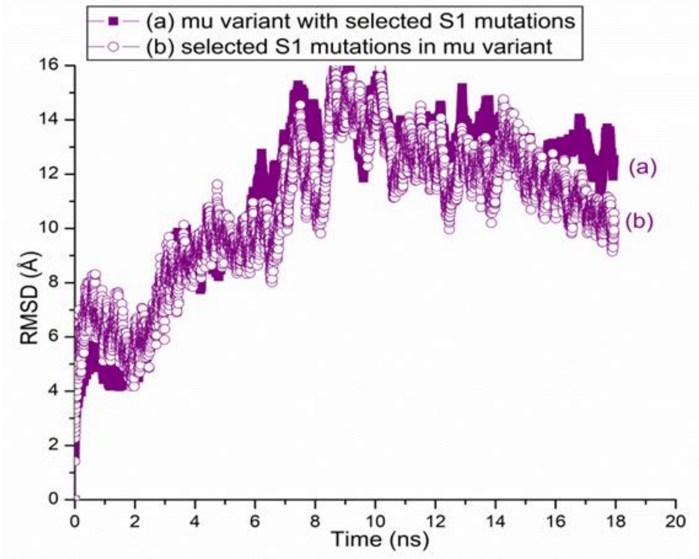


**Fig. S6.** The RMSD plots for the SARS-CoV-2 (a) Mu variant S1 and (b) S1 mutations within the Mu variant**.** The variant proteins are based on S1 model structure.

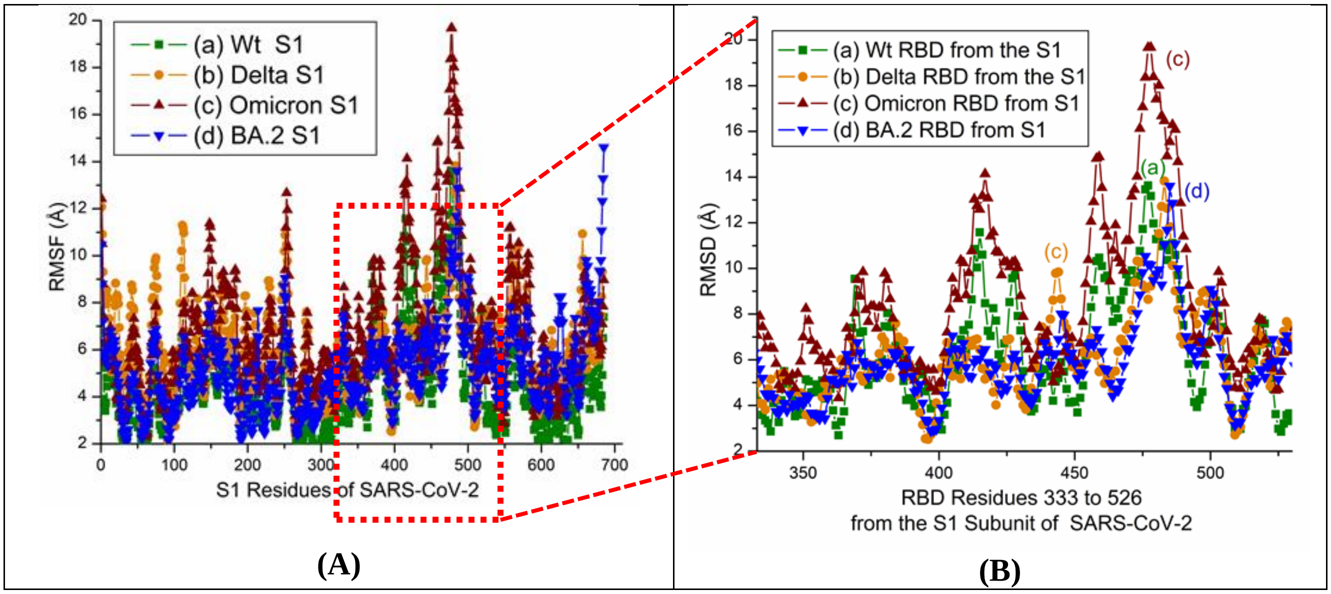


**Fig. S7.** RMSF plots of S1 in SARS-CoV-2 variants and subvariant. A. The RMSF plots of wt S1, Delta S, Omicron S1 and BA.2 S1 with selected mutations. B. Zoomed in view of the RBD residues 333 to 526 from the S1 Subunit of SARS-CoV-2 displayed in A. .

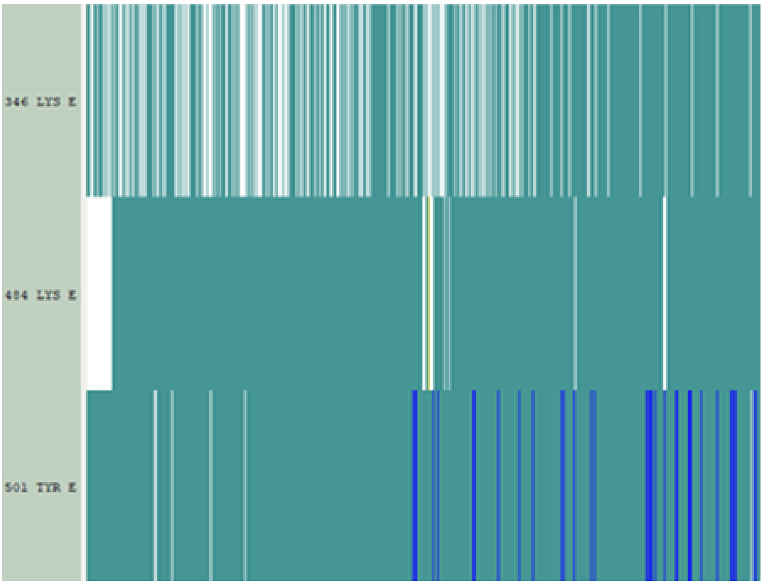


**Fig. S8.** Time based secondary structures of SARS-CoV-2 RBD with mutations R346K, E484K, and N501Y (observed predominantly in the Mu variant).

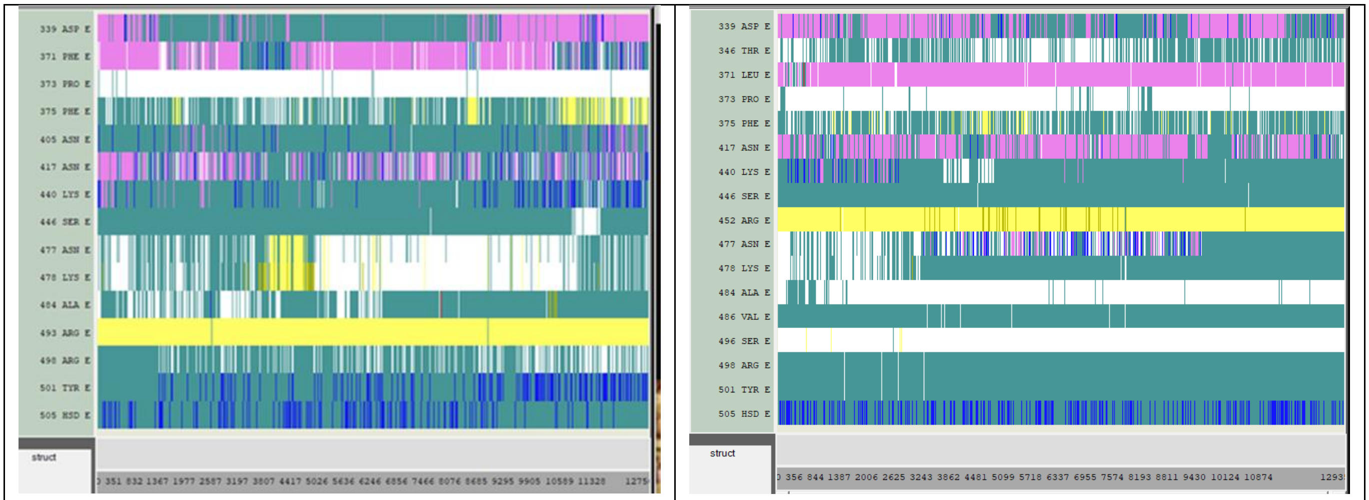


(A) (B)

**Fig. S9.** The time-based secondary structures of RBD mutations in (A) BA.2.75 and (B) BF.7

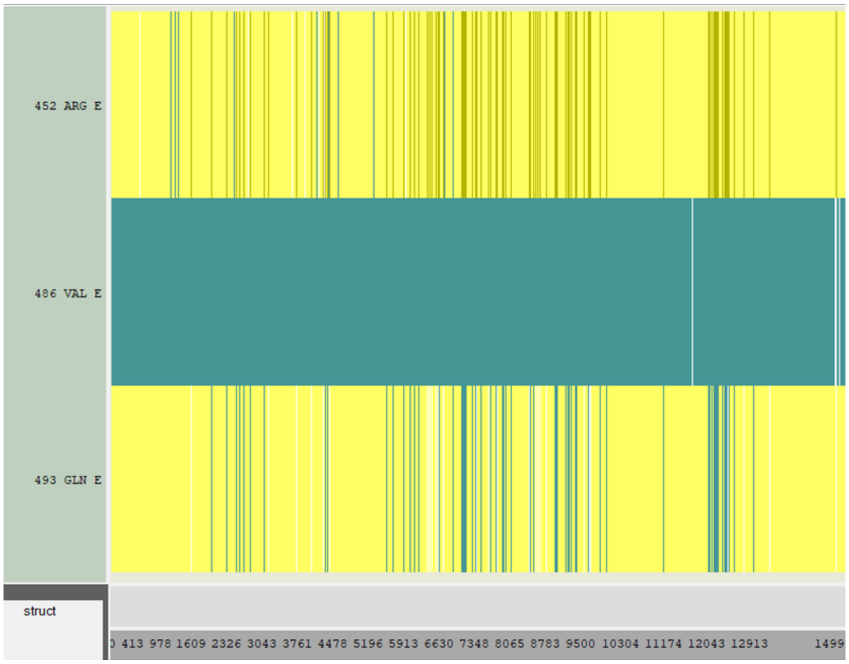


**Fig. S10.** Secondary RBD structure changes of a possible subvariant with mutations, L452R and F486V and reverse mutation Q493R. These two mutations and the reverse mutation are observed in subvariant BA.4/BA.5 in addition to the mutations that are observed in the sub variant BA.2.

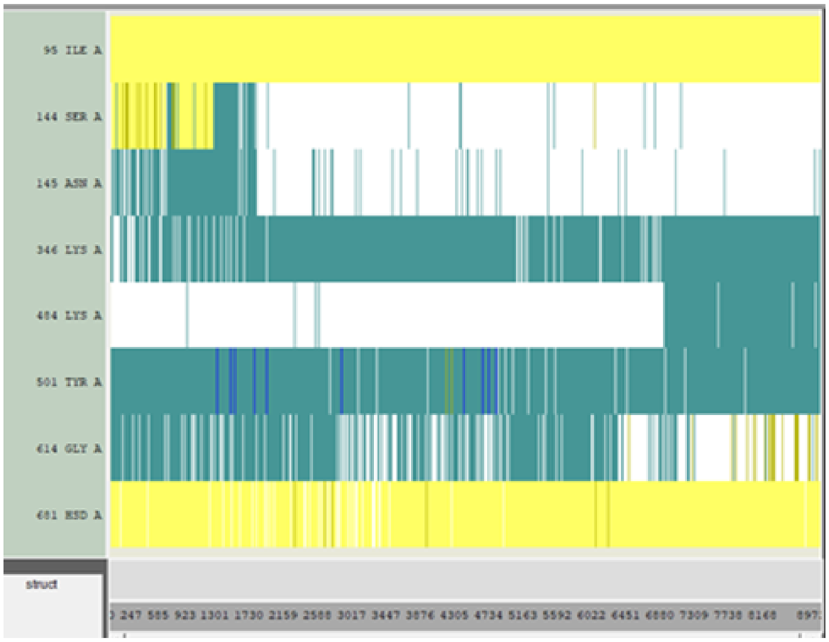


**Fig. S11**. Time based secondary structure changes of SARS-CoV-2 Mu variant with selected S1 mutations

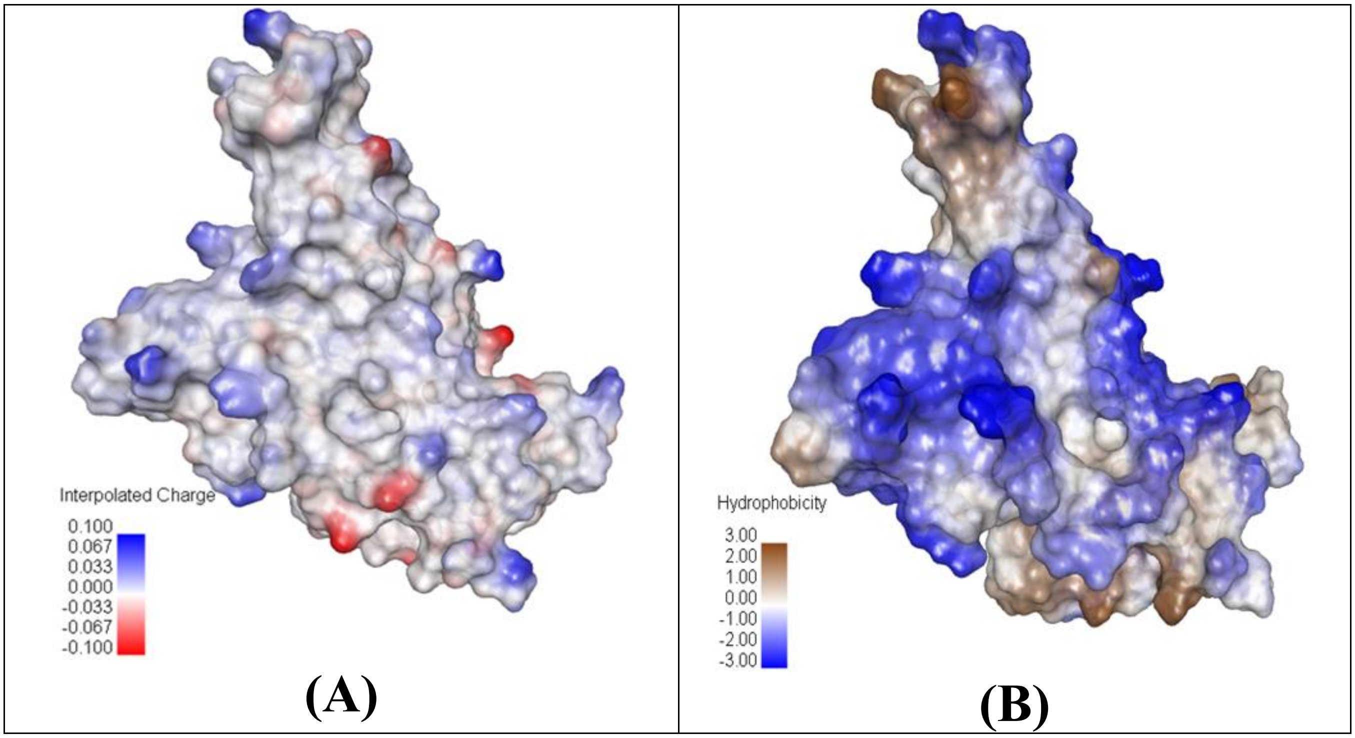


**Fig. S12.** A. Charged and B. Hydrophobic surface of BA.4/BA.5 RBD

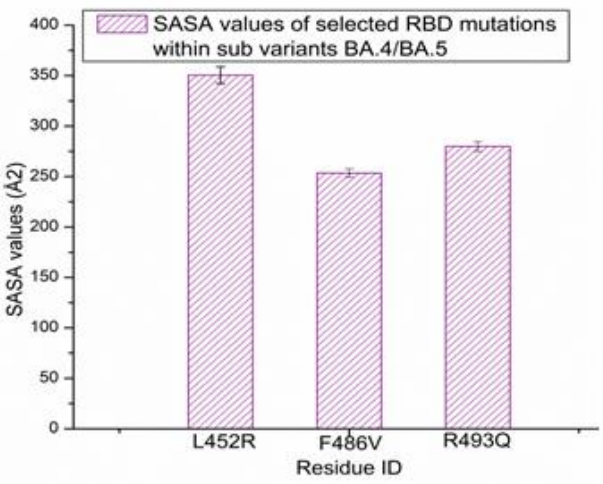


**Fig. S13.** SASA values of a possible subvariant with mutations, L452R and F486V and reversed mutation Q493R. These two mutations and the reversed mutation are observed in subvariant BA.4/BA.5 in addition to the mutations that are observed in the subvariant BA.2

### Table S1. Variants and subvariant of SARS-CoV2*

| Name of the lineage/variant | Reported mutations on the S protein | Set 1 Expt: simulation with key S1 RBD mutations | Set 2 Expt: simulation with key S1 mutations |
|---|---|---|---|
| Delta/ B.1.617.2 | T19R, V70F, T95I, G142D, del E156, del F157, R158G, A222V, W258L, K417N, L452R, T478K, D614G, P681R and D950N. | K417N, L452R and T478K. | T19R, V70F, T95I, G142D, R158G, A222V, W258L, K417N, L452R, T478K, D614G, and P681R. |
| Omicron/B.1.1.529 | A67V, del 69-70, T95I, del 142-144, Y145D, del211, L212I, ins214EPE, G339D, S371L, S373P, S375F, K417N, N440K, G446S, S477N, T478K, E484A, Q493R, G496S, Q498R, N501Y, Y505H, T547K, D614G, H655Y, N679K, P681H, N764K, D796Y, N856K, Q954H, N969K,.and 981F. | G339D, S371L, S373P, S375F, K417N, N440K, G446S, S477N, T478K, E484A, Q493R, G496S, Q498R, N501Y and Y505H. | A67V, T95I, Y145D, L212I, G339D, S371L, S373P, S375F, K417N, N440K, G446S, S477N, T478K, E484A, Q493R, G496S, Q498R, N501Y, Y505H, T547K, D614G, H655Y, N679K and P681H. |
| Omicron sub variant BA.2** | T19I, del 24-26, A27S, A67V, del 69-70, T95I, del 142-144, Y145D, del211, L212I, V213G, ins214EPE, G339D, S371F, S373P, S375F, T376A, D405N, R408S, K417N, N440K, G446S, S477N, T478K, E484A, Q493R, G496S, Q498R, N501Y, Y505H, T547K, D614G, H655Y, N679K, P681H, N764K, D796Y, N856K, Q954H, N969K and | G339D, S371F, S373P, S375F, T376A, D405N, R408S, , K417N, N440K, G446S, S477N, T478K, E484A, Q493R, G496S, Q498R, N501Y and Y505H. | T19I, A27S, A67V, T95I, Y145D, L212I, V213G, G339D, S371F, S373P, S375F, T376A, D405N, R408S, K417N, N440K, G446S, S477N, T478K, E484A, Q493R, G496S, Q498R, N501Y, Y505H, T547K, D614G, H655Y, N679K and |

| | | | |
|---|---|---|---|
| | L981F. | | P681H. |
| Mu/B.1.621 | T95I, Y144S, Y145N, R346K, E484K, N501Y, D614G, P681H and D950N. | R346K, E484K and N501Y. | T95I, Y144S, Y145N, R346K, E484K, N501Y, D614G and P681H. |

*A few mutations are observed in some of the sequences.
(Some of the mutations have been updated further through the course of this study).

** Recent report indicates that the Omicron subvariant BA.2 has multiple point mutations similar to those of Omicron variant. The BA.2 variant described in this report is based on older data.

### Table S2. Mutations in emerging subvariants of Omicron

| Name of the sub variant | Reported mutations on the S protein | simulation with key RBD mutations |
|---|---|---|
| BA.4/BA.5[+/++] | A67V, del69-70, T95I, del142-144, Y145D, del211, L212I, ins214EPE, G339D, S371L, S373P, S375F, K417N, N440K, G446S, L452R, S477N, T478K, E484A, F486V, G496S, Q498R, N501Y, Y505H, T547K, D614G, H655Y, N679K, P681H, N764K, D796Y, N856K, Q954H, N969K and L981F. | G339D, S371L, S373P, S375F, K417N, N440K, G446S, L452R, S477N, T478K, E484A, F486V, Q493R, G496S, Q498R, N501Y and Y505H. |
| BA.2.12.1[+] | G339D, S371F, S373P, S375F, T376A, D405N, R408S, K417N, N440K, L452Q, S477N, T478K, E484A, Q493R, Q498R, N501Y, Y505H and S704F. | G339D, S371F, S373P, S375F, T376A, D405N, R408S, K417N, N440K, L452Q, S477N, T478K, E484A, Q493R, Q498R, N501Y and Y505H. |
| BF.7[++] | A67V, del 69-70, T95I, del 142-144, Y145D, del211, L212I, ins214EPE, G339D, R346T, S371L, S373P, S375F, K417N, N440K, G446S, S477N, T478K, E484A, Q493R, G496S, Q498R, N501Y, Y505H, T547K, D614G, H655Y, N679K, P681H, N764K, D796Y, N856K, Q954H, N969K.and 981F. | G339D, R346T, S371L, S373P, S375F, K417N, N440K, G446S, L452R, S477N, T478K, E484A, F486V, G496S, Q498R, N501Y and Y505H. |
| BA.2.75[++/*] | W152R, F157L, I210V, G257S, D339H, G446S and N460K. | D339H, G446S and N460K |
| A possible variant with only additional two mutations that are observed in BA.4/ BA.5* | L452R and F486V. | L452R and F486V. |

[+] SARS-CoV-2 Variant Classifications and Definitions; https://www.cdc.gov/coronavirus/2019-ncov/variants/variant-classifications.html

++ https://www.who.int/en/activities/tracking-SARS-CoV-2-variants/
accessed June 14, 2022; August 18, 2022; 18 January 2023
(Some of the mutations have been updated further through the course of this study).

* SARS-CoV-2 variants of concern as of 9 June and 7 July 2022 https://www.ecdc.europa.eu/en/covid-19/variants-concern, accessed June 17, 2022; July 12, 2022,
\

### Table S3. RBD Mutations in circulating VOI and VUMs of SARS-CoV-2++

| Name of the sub variant | simulation with key RBD mutations |
|---|---|
| CH.1.1 | D339H, S371L, S373P, S375F, K417N, N440K, G446S, L452R, N460K, S477N, T478K, E484A, F486S, G496S, Q498R, N501Y and Y505H. |
| BQ.1./BQ1.1 | G339D, R346T, S371L, S373P, S375F, K417N, N440K, K444T G446S, N460K S477N, T478K, E484A, Q493R, G496S, Q498R, N501Y and Y505H. |
| XBB | G339H, R346T, L368I , S371L, S373P, S375F, K417N, N440K, V445P, G446S, N460K, S477N, T478K, E484A, F486S, F490S, Q493R, G496S, Q498R, N501Y and Y505H. |
| XBB1.5 | G339H, R346T, L368I , S371L, S373P, S375F, K417N, N440K, V445P, G446S, N460K, S477N, T478K, E484A, F486P, F490S, Q493R, G496S, Q498R, N501Y and Y505H. |
| XBF | G339H, R346T, S371L, S373P, S375F, K417N, N440K, G446S, N460K, S477N, T478K, E484A, F486P, F490S Q493R, G496S, Q498R, N501Y and Y505H. |

++ https://www.who.int/en/activities/tracking-SARS-CoV-2-variants/
Accessed 30 March 2023 (Some of the mutations have been updated further through the course of this study).
